# INDUCED MUTATIONS IN *TaASN-A2* REDUCE FREE ASPARAGINE CONCENTRATION IN THE WHEAT GRAIN

**DOI:** 10.1101/2021.11.08.467752

**Authors:** Rocío Alarcón-Reverte, Yucong Xie, John Stromberger, Jennifer D. Cotter, Richard Esten Mason, Stephen Pearce

**Affiliations:** Department of Soil and Crop Sciences, Colorado State University, Fort Collins, CO 80523, USA

## Abstract

Acrylamide is a neurotoxin and probable carcinogen formed as a processing contaminant during baking and production of different foodstuffs, including bread products. The amino acid asparagine is the limiting substrate in the Maillard reaction that produces acrylamide, so developing wheat varieties with low free asparagine concentrations in the grain is a promising approach to reduce dietary acrylamide exposure. A candidate gene approach was used to identify chemically-induced genetic variation in *ASPARAGINE SYNTHETASE 2* (*ASN2)* genes that exhibit a grain-specific expression profile. In field trials, durum and common wheat lines carrying *asn-a2* null alleles exhibited reductions in free asparagine concentration in their grains of between 9 and 34% compared to wild-type sister lines. These plants showed no significant differences in spikelet number, grain size and weight, germination or baking quality traits. These non-transgenic variants can be deployed without restriction in elite wheat germplasm to reduce acrylamide-forming potential with no negative impacts on quality or agronomic performance.

**Core ideas:**

- Three wheat *ASPARAGINE SYNTHETASE 2* knockout alleles were characterized in field experiments.
- Mutant alleles conferred significant reductions in grain free asparagine concentration.
- The alleles did not affect quality or agronomic traits.
- These non-transgenic alleles can be deployed without restriction in wheat breeding programs.

## INTRODUCTION

Acrylamide is a potent neurotoxin and causes cancer in rodents (Beland et al., 2013). For humans, acrylamide is classified as a probable carcinogen by the International Agency for Research on Cancer (IARC, 1994) and mutational signatures attributed to acrylamide exposure were detected in approximately one-third of assayed tumor genomes, including 19 different tumor tissues (Zhivagui et al., 2019). There is a strong and growing incentive to reduce potential sources of acrylamide exposure in humans, and regulatory agencies are beginning to specify limits in recognition of this threat (Reviewed in (Raffan & Halford, 2019)).

A major source of acrylamide exposure is the consumption of processed foods that are rich in carbohydrates (Tareke et al., 2002). Acrylamide levels are highest in potatoes and coffee, but wheat (*Triticum aestivum* L.) is also a major source of dietary acrylamide due to the high volume of bread products consumed in the human diet (Raffan & Halford, 2019). Acrylamide is a processing contaminant that accumulates in the high temperature and low moisture conditions during baking as a product of the Maillard reaction (Mottram et al., 2002; Stadler et al., 2002). Modifying production conditions such as baking at lower temperatures or adding chemical amendments can reduce the level of acrylamide formation, but these are often impractical to implement (Claus et al., 2008). Because asparagine provides the carbon skeleton on which acrylamide is formed, free asparagine concentration is the limiting substrate for acrylamide formation (Surdyk et al., 2004) and, in wheat, free asparagine concentrations in the grain are highly correlated with acrylamide levels in baked products (R^2^ > 0.99) (Curtis et al., 2014). Therefore, one promising approach to reduce dietary exposure to acrylamide is to develop wheat varieties with low free asparagine concentration in grains.

Free asparagine concentration is strongly influenced by environmental factors. Providing nitrogen is abundant, asparagine accumulates in the grain when protein synthesis is limited, such as in response to biotic and abiotic stresses, including Sulphur (S) deficiency (Lea et al., 2007). Therefore, growers can minimize free asparagine concentrations by keeping crops free from pathogens and ensuring sufficient S supply in the soil (Claus et al., 2006; Curtis et al., 2019; Wilson et al., 2020). The estimated heritability of free asparagine concentration ranges between 0.13 to 0.65 in different field studies and exhibits approximately three- to five-fold variation within different panels of wheat germplasm (Corol et al., 2016; Emebiri, 2014; Rapp et al., 2018). Free asparagine concentration is a costly trait to phenotype, so characterizing the genetic basis of this variation would allow breeders to apply marker assisted or genomic selection to lower free asparagine concentration. However, the trait is highly quantitative with a complex genetic architecture influenced by multiple small-effect Quantitative Trait Loci (QTL) (Emebiri, 2014; Navrotskyi et al., 2018; Rapp et al., 2018). Few of these QTL overlap between studies, likely a factor of the strong influence of genotype by environment (G × E) interactions.

A complementary strategy to reduce free asparagine concentrations is to leverage knowledge of asparagine biosynthesis to target candidate genes. In eukaryotes, asparagine synthetase enzymes catalyze the Adenosine Triphosphate (ATP)-dependent assimilation of inorganic nitrogen in the form of ammonium into asparagine (Gaufichon et al., 2010). The wheat genome contains three homeologous copies of five asparagine synthetase genes that exhibit distinct expression profiles during development (Curtis et al., 2019; Oddy et al., 2021; Xu et al., 2017). The *TaASN2* genes are notable for their grain-specific expression profile, and a natural deletion of *TaASN-B2* is associated with a mild reduction in free asparagine concentrations in the grain (Oddy et al., 2021). By contrast, the *TaASN-A2* and *TaASN-D2* homeologous genes show higher transcript levels than *TaASN-B2* but exhibit no natural genetic variation among a collection of 14 wheat varieties (Oddy et al., 2021). In addition, no QTL for asparagine concentration has been mapped within the vicinity of an asparagine synthetase gene (Emebiri, 2014; Rapp et al., 2018). Plants carrying Clustered Regularly Interspaced Short Palindromic Repeats/CRISPR-associated protein (CRISPR/Cas)-induced knockouts of all three *TaASN2* homeologues exhibit reductions in free asparagine concentrations of up to 90% (Raffan et al., 2021). In greenhouse trials, these plants showed no obvious reduction in performance except a lower germination rate (Raffan et al., 2021). However, field trials are often necessary to fully characterize the impact of a genetic variant. For example, potato lines carrying ribonucleic acid interference (RNAi) constructs to suppress *StASN1* and *StASN2* transcripts exhibited significant reductions in tuber acrylamide levels compared to wild-type control lines in the greenhouse (Rommens et al., 2008), but subsequent field trials revealed these tubers were smaller and had a cracked appearance (Chawla et al., 2012). A more targeted approach using an RNAi construct specific to *StASN1* controlled by a tuber-specific promoter had a similar impact in reducing tuber asparagine concentrations without impacting tuber yield in field-grown plants (Chawla et al., 2012).

For wheat, it will be important to evaluate the association between asparagine concentration and quality traits (Claus et al., 2006). Asparagine concentration showed a mild negative correlation with sedimentation volume (Z-sodium dodecyl sulfate, Z-SDS) (Corol et al., 2016; Rapp et al., 2018), a measure of the gluten quantity and strength in a flour sample. In a panel of European wheats, free asparagine concentration was also negatively correlated with thousand grain weight (TGW) and hectoliter volume, and positively correlated with protein content (Rapp et al., 2018). A range of quality tests have now been optimized to more efficiently assay quality traits from small volumes of wheat grain. The Single Kernel Characterization System (SKCS) measures grain hardness, moisture, diameter and weight (Gaines et al., 1996), the solvent retention capacity (SRC) assay is an indicator of gluten strength, gliadins, pentosans and starch damage (Bettge et al., 2002), while mixograph analyses provide an indication of gluten strength and dough rheological properties (Bloksma & Bushuk, 1988).

In the current study, three backcross populations segregating for Ethyl-MethaneSulfonate (EMS)-induced null alleles of *ASN-A2* were developed in durum and common wheat. In field trials, mutant lines exhibited significant reductions in free asparagine concentrations in their grain compared to wild-type sister lines but showed no significant changes in agronomic or quality traits. These non-transgenic *asn-a2* null alleles can be utilized without restriction in breeding programs to help develop wheat cultivars with reduced acrylamide-forming potential.

## MATERIALS AND METHODS

### Plant materials

EMS-mutagenized M_4_ lines carrying null alleles in *ASN-A2* from the common wheat (*Triticum aestivum* L.) variety ‘Cadenza’ (T6-1048) and the durum wheat (*Triticum turgidum* L. subsp. *durum* (Desf.) Husn.) variety ‘Kronos’ (T4-2032 and T4-1388) were identified from an *in silico* TILLING database (Krasileva et al., 2017) and provided by Dr. Jorge Dubcovsky, UC Davis. Each line was backcrossed twice to non-mutagenized wild-type plants of the corresponding variety (‘Kronos’ for lines T4-2032 and T4-1388; ‘Cadenza’ for line T6-1048) to reduce the impact of background mutations. In 2019, selections were made of BC_1_F_2_ homozygous wild-type and *asn-a2* sister lines to grow in field experiments. In 2020, similar selections were made of homozygous wild-type and *asn-a2* sister lines using BC_2_F_2_ materials. These sister lines were grown in greenhouse conditions for qRT-PCR experiments to quantify

### ASN2 transcript levels

Seeds of the winter wheat varieties Antero, Hatcher, Snowmass, Snowmass 2.0, Brawl, Byrd, Denali, SY Wolf, Winterhawk and Canvas were sown in field experiments in 2017 and 2018 and were sourced from the CSU wheat breeding program from Dr. Scott Haley.

### Field experiments

In the 2016-2017 field season, free asparagine concentration was measured in the grains of eight elite winter wheat varieties (Antero, Hatcher, Snowmass, Brawl, Byrd, Denali, SY Wolf and Winterhawk) harvested from field plots in five Colorado State University wheat breeding program trial location (Fort Collins 40.653086 -104.999863, Akron 40.14858333, -103.1395472, Burlington 39.2999083, -102.2983472, Julesburg 40.8996, -102.2293278, Yuma 40.18595, - 102.66099). In the 2017-2018 field season, asparagine concentration was measured in four elite winter wheat varieties (Snowmass, Snowmass 2.0, Byrd, Canvas) in seven locations (Fort Collins, Akron, Julesburg, Orchard 40.479692, -104.071244, New Raymer 40.56793056, - 103.8940111, Yuma, and Roggen 40.088177, -104.261670).

In the 2018-2019 field season, BC_1_F_2_ sister lines homozygous for wild-type and mutant alleles of T4-2032, T4-1388 and T6-1048 were grown as headrows in Fort Collins using a randomized complete block design. Six genotypes were placed randomly within each block with five replications. Due to the loss of some samples some genotypes had fewer replicates. Free asparagine concentration was measured from the grains of these lines. This experiment was repeated in the 2019-2020 field season in the same location using a randomized complete block design with six genotypes and six replications using BC_2_F_2_ sister lines and were phenotyped for a broader range of traits, including free asparagine content in grain, mean kernel weight (mg), spikelets per spike, germination rate five, seven and 14 days after sowing, seed diameter (mm), hardness index, protein concentration in grain, grain moisture (%) and solvent retention capacity of the flour.

### Phenotyping

Spikelet number was measured as the average number from ten randomly-selected spikes from each headrow. Germination tests were carried out using 50 seeds from each of the 36 headrows from the field experiment from field season 2019-2020. Seeds from each headrow were placed in two pots filled with compost from 24 cell seed inserts (25 seeds each) organized in five lines of five seeds each. Pots were placed in standard 1020 seeds trays in the greenhouse with these conditions: 19.4 °C average day temperature, 18.3 °C average night temperature, 40% RH and 16 h daylength. Emerged seedlings were counted in each pot five, seven and 14 days after sowing.

Quality evaluations were run at CSU’s wheat quality laboratory using approved methods from the American Association of Cereal Chemists (AACC, 2000). The Single Kernel Characterization System (SKCS 4100; Perten Instruments) was used to measure mean kernel weight (mg), diameter (mm), and hardness index.

Protein concentration was measured by near-infrared spectroscopy (NIRS) on whole-grain and wheat meal samples using a Foss NIRSystems model DA1650 (Foss North America Inc.). A validation set of 25 samples were analyzed using the combustion method and used to adjust NIR calibrations if necessary (Leco Model FP-428; Approved Method 46-30). Samples of grain were ground on a UDY mill (UDY Corporation, Fort Collins, CO) equipped with a .5 mm sieve. Grain protein concentrations are presented as 12% moisture basis, and flour protein concentrations on a 14% moisture basis. Dough-mixing properties were assayed using a 10 g mixograph according to AACC Method 54-40A, which was optimized for water absorption based on flour protein concentration obtained from the NIRS according to the equation:

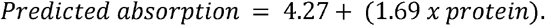

where protein represents the flour protein concentration (obtained with NIRS) at 140 g kg^−1^ moisture basis. The prediction equation was determined from experiments with the hard red winter wheat cultivar ‘Cheyenne’ (PI 553248; B. Seabourn, personal communication, 2003).

Dough-mixing properties were determined using the computerized mixograph and MIXSMART software (v. 1.0.484 for Windows, National Manufacturing). The following computerized mixograph parameters were used in the analyses: midline peak time, midline peak height, midline peak width, midline right slope, and midline right width (the midline curve width two min after peak).

Solvent retention capacity (SRC) analysis was performed using whole wheat flour for water, sucrose, lactic acid and carbonate solvents, using a method described previously (Bettge et al., 2002).

### Asparagine quantification

#### Sample preparation

Following harvest, 2-3 g of seeds form each plot were milled using a hand mill and 95-105 mg of whole flour were placed in 2 mL Eppendorf tubes for asparagine quantification. Homogenized flour samples (∼100 mg) were accurately weighed and mixed with 1 mL of cold 80% methanol in water. 50 µg of isotopically labeled 13C4 L-asparagine (99%, Cambridge Isotope Laboratories, MA) as internal standard (IS) were added from 1 mg/mL stock solution to each sample. Samples were vigorously vortexed at 4°C for 1 h, and then sonicated for 30 min before an additional 1-h vortexing at 4°C followed by centrifugation at 3,000 rpm for 15 min at 4°C. 750 µL of supernatant were recovered and solvent was removed under nitrogen. To the dried extract, 600 µL of methyl-tert-butyl ether (MTBE), 300 µL of methanol and 100 µL of water were added. Samples were vigorously vortexed for 1 min, and then 500 µL of water were added to induce phase separation. The upper organic layer containing unwanted lipids was removed and discarded. To ensure all lipids were removed, another 1 mL of MTBE was added to the samples, vortexed for 1 min, centrifuged again at 3,000 rpm for 10 min and the upper organic layer was again discarded. The solvent in lower layer was removed under nitrogen. The dried samples were re-suspended in 300 µL of water before the addition of 700 µL of acetonitrile and vortexed for 1 min and then incubated at 4°C for 1 h before centrifugation at 17,000 g and 4°C. 900 µL of the upper supernatant were recovered, dried under nitrogen gas, re-suspended in 50 µL of 50% methanol in water (v/v) and stored at -20 °C until analysis. Samples for a calibration curve for quantification were prepared in 50% methanol in water (v/v) at concentrations ranging from 0-200 µg/mL with IS and were analyzed in triplicates.

#### Liquid Chromatography-Mass Spectrometry Analysis

Each sample (2 µL) was injected onto a Waters Acquity Ultra-performance liquid chromatography (UPLC) system in a randomized order with a pooled quality control (QC) injection after every six sample injections and separated using a Waters Acquity UPLC ethylene bridged hybrid Amide column (1.7 µM, 2.1 × 30 mm) at a constant flow rate of 0.4 µL/min. The elution solvents were: A, acetonitrile with 0.1% formic acid; B, water with 10 mM ammonium formate and 0.1% formic acid. The solvent gradient started with 0.1% B at 0 min, increased to 80% B at 1.5 min, held at 80% B until 2.0 min, re-equilibrated to 0.1% B at 2.25 min, held at 0.1% B for 2.5 minutes. The total run time was 4.75 min. The column and samples were held at 45 °C and 4 °C, respectively. The column eluent was infused into a Waters Xevo G2 TOF-MS with an electrospray source in positive mass spectrometry (MS) fullscan mode with target enhancement at m/z 133. The dwell time was 0.2 s. The collision energy was set at 0. Calibration was performed using sodium iodide with 1 ppm mass accuracy. The capillary voltage was held at 2200 V, source temperature at 150 °C, and nitrogen desolvation temperature at 350 °C with a flow rate of 800 L/hr. The ions used for quantification are listed below.

* IS, internal standard

|  | Quantification, [M+H]<sup>+</sup>, *m/z* | Confirmation, *m/z* | Confirmation, *m/z* |
| --- | --- | --- | --- |
| Asparagine | 133.02 | 87.05 | 74.02 |
| IS* | 137.03 | 90.06 | 76.03 |

### Quality Control

QC samples were pooled from final extracts of all the actual samples and were injected after every 6 actual samples. Mean values and standard deviations for QC can be found in the result sheet. The CV of 8 QCs was 8.4%.

#### Statistical analysis

ANOVA single factor analysis using the Data Analysis package from Excel was performed to calculate statistical significance between genotypes in individual lines. To calculate statistical significance of genotypes versus lines an ANOVA Type III analysis was performed using the computer software JASP (JASP, 2019).

#### Genotyping assays

Genomic DNA was extracted from seedling leaf tissue using the Cetyl TrimethylAmmonium Bromide (CTAB) method (Murray & Thompson, 1980). Kompetitive Allele Specific PCR (KASP) assays were developed to genotype the G468A mutation in line T4-1388 (Supplemental Figure S1) and the G446A mutation in line T4-2032 (Supplemental Figure S2). Primers for each assay are listed in Supplemental Table S1. Each reaction consisted of 0.14 µL of KASP primer mix, 5 µL of KASP Master mix (LGC Genomics, LLC, Beverly, MA, USA) and 5 µl of template DNA (50 ng/µl). A 100 µL KASP primer mix stock was previously prepared using 46 μL nuclease-free water, 30 μL common primer (100 μM), 12 μL specific primer with FAM (100 μM) and 12 μL specific primer with VIC (100 μM). The KASP thermal cycling conditions were: 94 □ for 15 s; 10 cycles of: 94 □ for 15 s, 61 □ for 1 min; 26 cycles of: 94 □ for 20 s, 55 □ for 1 min; 30 □for 1 min.

The G585A mutation in line T6-1048 was genotyped using a Cleaved Amplified Polymorphic Sequences (CAPS) marker (Supplemental Figure S3). Primers ASN-A2_1048_F3 and ASN-A2_1048_R3 (Supplemental Table S1) were used to amplify a 1,063 bp product using the conditions 95 □ for 30 s; 35 cycles of: 95 □ for 15 s, 53 □ for 30 s, 68 □ for 1 min; 68 □ for 5 min. Each reaction consisted of 2.5 µL 10X Standard *Taq* Reaction Buffer (NEB, Ipswich, MA, USA), 0.5 µL 10 mM dNTPs (Invitrogen, Life Technologies, Carlsbad, CA, 92008, USA), 0.5 µL 10 µM Forward Primer, 0.5 µL 10 µM Reverse Primer, 5 µL Template DNA (50 ng/µl), 0.125 µL *Taq* DNA Polymerase (NEB, Ipswich, MA, USA) and nuclease-free water to a total reaction volume of 25 µL.

Amplified DNA was digested at 37 L for 2 h with the restriction enzyme *Sty*I-HF (NEB, Ipswich, MA, USA) and products were run on a 0.8 % agarose gel stained with Invitrogen™ SYBR™ Safe DNA Gel Stain (Life Technologies, Carlsbad, CA, USA). Template DNA carrying the mutant allele (A at nucleotide 585) showed a 1,063 bp band, while the wild-type allele (G at nucleotide 585) showed two fragments of 946 bp and 117 bp (Supplemental Figure S3). Heterozygous individuals had a characteristic banding pattern of 1,063 bp, 946 bp and 117 bp. The presence and absence of *ASN-B2* was determined using an assay described previously (Oddy et al., 2021).

#### Quantitative Reverse Transcriptase PCR (qRT-PCR)

Developing grains from six biological replicates of each genotype were harvested at two time points (21-, and 28-days post anthesis) and immediately frozen in liquid nitrogen. Grains were ground to a fine powder using a pestle and mortar and RNA was extracted using a standardized extraction protocol modified from the CTAB method (Chang et al., 1993). RNA was quantified using a NanoDropTM 1000 Spectrophotometer (Thermo Fisher Scientific). To remove genomic DNA contamination, 1 µg RNA was treated with SuperScriptTM IV VILO ezDNase Enzyme (Thermo Fisher Scientific) in a 10 µL solution and incubated for 4 min at 37 □.

The DNase-treated RNA samples were quantified using a NanoDropTM 1000 Spectrophotometer before cDNA synthesis. cDNA was synthesized by mixing the ezDNase enzyme treated RNA samples in nuclease-free water with SuperScriptTM IV VILO Master Mix (Invitrogen), according to the manufacturer’s instructions, and placing in a thermocycler using the following program: 25 □ for 10 min; 50 □ for 30 min; 85 □ for 5 min. Samples were diluted to 10 ng/µL according to the RNA-equivalent of starting DNAse-treated sample before synthesis.

qRT-PCR was performed using the QuantStudioTM 3 System (Applied Biosystems) set to standard curve mode in 96-well 0.1-mL block. Each reaction contained 10 µL SYBR Green Master Mix (Applied Biosystems), 1 µL forward and reverse primer (10 mM) and 5 µL cDNA (10 ng/µL). The expression of each target gene was measured as 2^(-Δ_CT) relative to *ACTIN*. Primers for *ASN-A2, ASN-B2* and *ASN-D2* were described previously (Oddy et al., 2021). The qRT-PCR primers for *ASN-A2* amplify a region of the gene in exon 1, upstream of all the induced EMS mutations used in the materials of this study.

Graphs were plotted in R using ggplot2 and student t-tests were performed to calculate statistical significance of pair-wise differences in expression between genotypes.

## RESULTS

### Characterization of asn-a2 null alleles

Two EMS-mutagenized lines in tetraploid ‘Kronos’ (T4-1388 and T4-2032) and one in hexaploid ‘Cadenza’ genetic backgrounds (T6-1048) were identified that carry point mutations predicted to introduce premature stop codons in the *ASN-A2* coding region (Figure 1a). All three lines are predicted to encode C-terminally truncated ASN-A2 proteins lacking the entire ASN-synthetase domain (Figure 1b), so are highly likely to encode non-functional proteins. Both ‘Kronos’ and ‘Cadenza’ carry a complete *ASN-B2* gene (Supplemental Figure S4), so T4-1388 and T4-2032 mutant plants carry one functional *ASN2* gene (*TdASN-B2*) and T6-1048 mutant plants carry two functional *ASN2* genes (*TaASN-B2* and *TaASN-D2*). Segregating BC_1_F_2_ and BC_2_F_2_ populations for each of these null alleles were developed by backcrossing to the corresponding wild-type parental line, selecting mutant alleles using genotyping assays described Supplemental Figures S1, S2 and S3.

**Figure 1:**
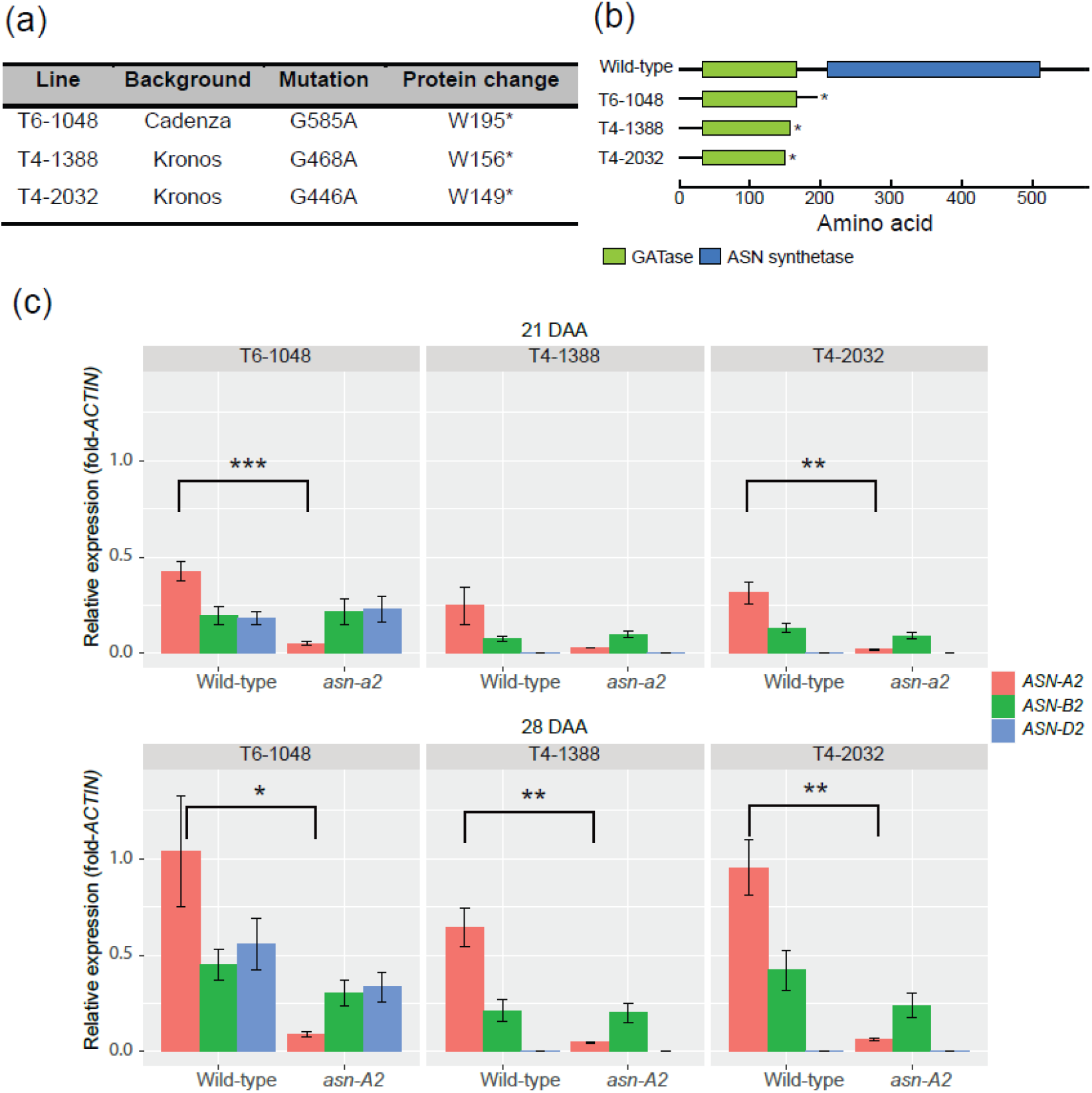
*ASN-A2* mutant characterization. **(a)** Details of the induced mutations in each line. Mutation position represents the mutated residue in the wild-type *ASN-A2* coding sequence. Protein mutations refer to the amino acid mutated where W = Tryptophan and * = stop codon. (**b)** Schematic representation of the selected mutations in each line and their effects on protein translation. Glutamine amidotransferase (GATase) and ASN synthase conserved domains are highlighted in green and blue, respectively, and drawn to scale based on protein size. (**c)** Transcript levels of *ASN2* genes in grain tissues of wild-type and mutant lines at 21 days after anthesis (DAA) and 28 DAA. Error bars represent standard error of the mean (n = 6, except for T4-2032 wild-type samples, where n = 5). Student t-tests were performed for each gene between genotypes at each timepoint, * = *P* < 0.05; ** = *P* < 0.01; *** *P* < 0.001.

In wild-type plants of all three lines, *ASN-A2* was the most highly expressed homeologue in grain tissues, and transcript levels rose between 21 days after anthesis (DAA) and 28 DAA (Figure 1c). In all three mutant genotypes, *ASN-A2* transcript levels were significantly lower than in wild-type sister lines at 28 DAA, and for T6-1048 and T4-2032, were also significantly lower at 21 DAA (Figure 1c). By contrast, *ASN-B2* and *ASN-D2* transcript levels were lower and not differentially expressed between wild-type and mutant genotypes at either timepoint (Figure 1c).

### Effect of asn-a2 mutations on free asparagine concentration

To assay the effect of the *asn-a2* mutations on free asparagine concentrations, mature grain was harvested from field-grown headrows of BC_1_F_2_ sister lines from each population as five biological replicates. Free asparagine concentration was significantly lower in *asn-a2* mutants compared to wild-type sister lines in the populations derived from line T6-1048 (29% reduction, *P* < 0.001) and line T4-1388 (33% reduction, *P* < 0.05, Figure 2a). In the population derived from line T4-2032, asparagine concentration was 9% lower in the *asn-a2* mutant plants, but the difference was not significant (*P* = 0.198, Figure 2a). These results were consistent with BC_2_F_2_ populations grown in 2020 at the same location as headrows in six biological replicates (Figure 2b). Free asparagine concentration was significantly lower in *asn-a2* mutants compared to wild-type sister lines in the populations derived from line T6-1048 (28% reduction, *P* < 0.0001) and line T4-1388 (24% reduction, *P* < 0.05). In line T4-2032, although the reduction in asparagine concentration in *asn-a2* mutant plants was proportionally greater than in the other populations (34% reduction) these differences were not significant due to higher variation (*P* = 0. 071, Figure 2b). Across both years of the experiment, genotype was highly significantly associated with free asparagine concentration (*P* < 0.0001).

**Figure 2:**
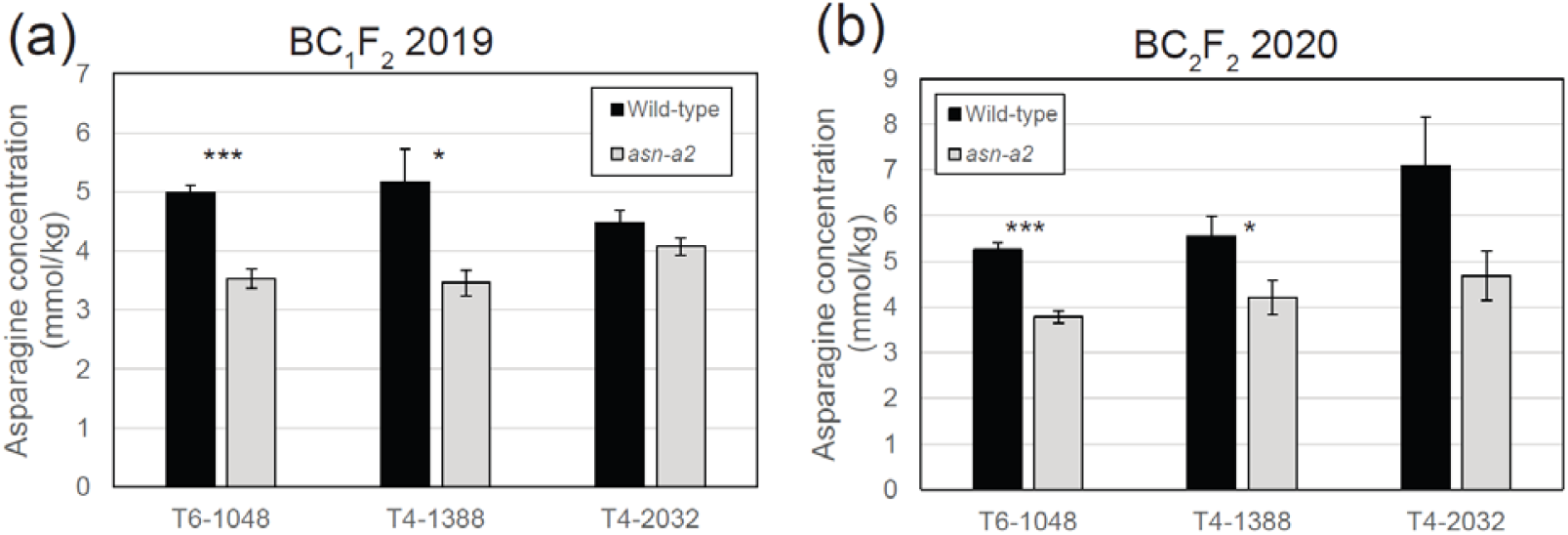
Free asparagine concentration in mature grain of wild-type and mutant *asn-a2* sister lines in (**a)** BC_1_F_2_ materials grown in 2019 and (**b)** BC_2_F_2_ materials grown in 2020. * = *P* < 0.05; *** *P* < 0.001.

### Effect of asn-a2 mutations on agronomic traits

To determine the impact of the *asn-a2* mutations on plant growth and development, agronomic traits were analyzed in grains harvested from field-grown BC_2_F_2_ plants. Germination rates were above 84% in all samples and there were no significant differences between wild-type and *asn-a2* genotypes, measured 5, 7 and 14 days after sowing (Table 1). In line T6-1048, the germination rate was between 2.7% and 3% lower in *asn-a2* mutants compared to wild-type sister lines, although the differences were not significant (Table 1). Conversely, germination rate was slightly higher in the *asn-a2* mutant compared to wild-type sister lines in T4-1388 and T4-2032 populations (Table 1). Despite variation in spikelet number, grain weight and diameter between ‘Cadenza’ and ‘Kronos’ genotypes, there were no significant differences in these traits between wild-type and *asn-a2* mutant sister plants in any of the three lines (Table 1).

**Table 1:** Effect of *asn-a2* mutations on agronomic traits in three lines. Genotype effect represents significance of genotype across all lines. * To measure diameter in T4-2032, five biological replicates were used.

| Trait | T6-1048 (n=6) |  |  | T4-1388 (n=6) |  |  | T4-2032 (n=6)* |  |  | Genotype effect |
| --- | --- | --- | --- | --- | --- | --- | --- | --- | --- | --- |
|  | Wild-type | *asn-a2* | *P* | Wild-type | *asn-a2* | *P* | Wild-type | *asn-a2* | *P* | *P* |
| Germination 5d | 89.0±1.612 | 86.3±1.202 | 0.214 | 88.3±1.745 | 89.3±0.989 | 0.629 | 84.7±1.606 | 87.0±1.612 | 0.329 | 0.862 |
| Germination 7d | 92.0±1.789 | 89.0±1.238 | 0.198 | 91.0±1.693 | 94.0±1.265 | 0.186 | 86.0±1.789 | 91.0±1.693 | 0.070 | 0.264 |
| Germination 14d | 92.7±1.606 | 89.7±1.085 | 0.153 | 92.7±1.605 | 94.7±0.667 | 0.277 | 86.3±1.892 | 91.7±1.498 | 0.052 | 0.315 |
| Spikelet number | 20.6±0.138 | 20.2±0.138 | 0.131 | 15.1±0.249 | 14.9±0.218 | 0.660 | 14.9±0.259 | 14.9±0.123 | 1 | 0.301 |
| Weight (mg) | 24.9±1.200 | 26.4±0.728 | 0.324 | 33.2±0.779 | 32.8±1.194 | 0.772 | 33.8±0.802 | 33.8±0.944 | 0.992 | 0.667 |
| Diameter (mm) | 2.50±0.048 | 2.50±0.27 | 0.439 | 2.80±0.038 | 2.80±0.056 | 0.956 | 2.84±0.040 | 2.83±0.047 | 0.870 | 0.727 |

### Effect of asn-a2 mutations on quality and breadmaking traits

The impact of the *asn-a2* null alleles on quality traits were tested using the Single Kernel Characterization System (SKCS) and NIR protein analysis, which revealed there were no significant differences between wild-type and *asn-a2* sister lines for grain moisture, hardness, grain protein concentration, flour protein concentration or flour ash concentration (Table 2). Solvent Retention Capacity (SRC) analysis revealed no significant differences between wild-type and mutant genotypes in lactic acid, carbonate, water or sucrose (Table 2), indicating these mutations have no impact on pentosan or gliadin content in the grains, or on starch and gluten quality.

**Table 2:**
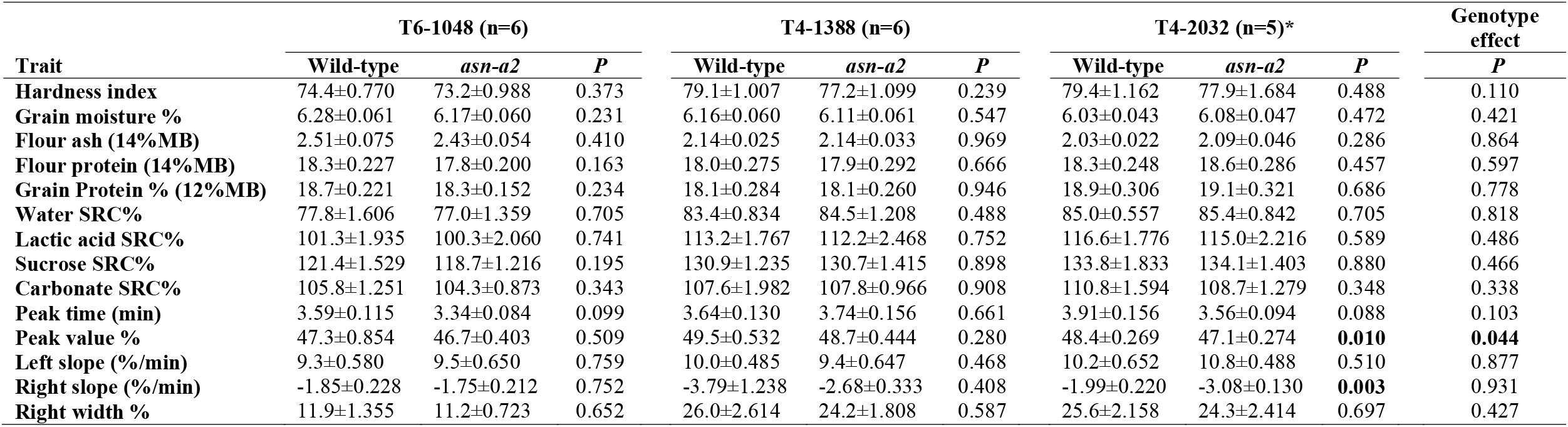
Effect of *asn-a2* mutations on quality traits. Significant effects are highlighted in bold. Genotype effect represents significance of genotype across all lines. SRC = Solvent Retention Capacity.

A mixograph was run on each sample to measure dough rheological properties and showed that *asn-a2* and wild-type sister lines exhibited no significant differences for peak mixing time, a measure of dough strength (Table 2). Peak value % was lower in *asn-a2* mutants than wild-type sister plants in all three lines, although this different was significant only in line T4-2032 (Table 2). Right slope (%/min) values were inconsistent between lines and were slightly higher (less negative) in *asn-a2* mutants in lines T6-1048 and T4-1388, but significantly lower (more negative) in line T4-2032 (Table 2). Right-width % was not significantly different between genotypes in any line, indicating that the *asn-a2* mutations do not confer differences in mixing tolerance. Full mixographs for all lines are provided in Supplemental Figure S5.

### Free asparagine concentration in wheat varieties adapted to the Great Plains

To better understand the opportunities to reduce acrylamide-forming potential in breeding programs, free asparagine concentration was measured in grain samples of elite winter wheat germplasm selected for their economic importance and acreage in Colorado and the Great Plains. Six winter wheat varieties were assayed in five field locations in 2017 (Figure 3a) and four varieties were assayed in seven locations in 2018 (Figure 3b). Asparagine concentrations varied by environment, with higher values in varieties grown in Fort Collins 2017 compared to Julesberg or Yuma (Figure 3a), but comparatively smaller differences between these environments in 2018 (Figure 3b). Some genotypes showed consistent free asparagine concentrations between environments. In 2017, ‘SY Wolf’ consistently exhibited high asparagine concentration in different environments, whereas in 2018, Snowmass exhibited the highest free asparagine concentration of all assayed varieties (Figure 3b). The varieties ‘Snowmass’, ‘Antero’ and ‘Hatcher’ carry the *TaASN-B2* deletion, while ‘Brawl’, ‘Byrd’, ‘Denali’, ‘SY Wolf’ and ‘Winterhawk’ all carry a functional copy of this gene (Supplemental Figure S4). In neither year was there a correlation between the presence or absence of *TaASN-B2* and mean asparagine concentration, indicating that other genotypic and environmental factors are driving the observed variation. The introgression of the *asn-a2* null allele into ‘Snowmass 2.0’ has been initiated to characterize its effect in an elite wheat variety.

**Figure 3:**
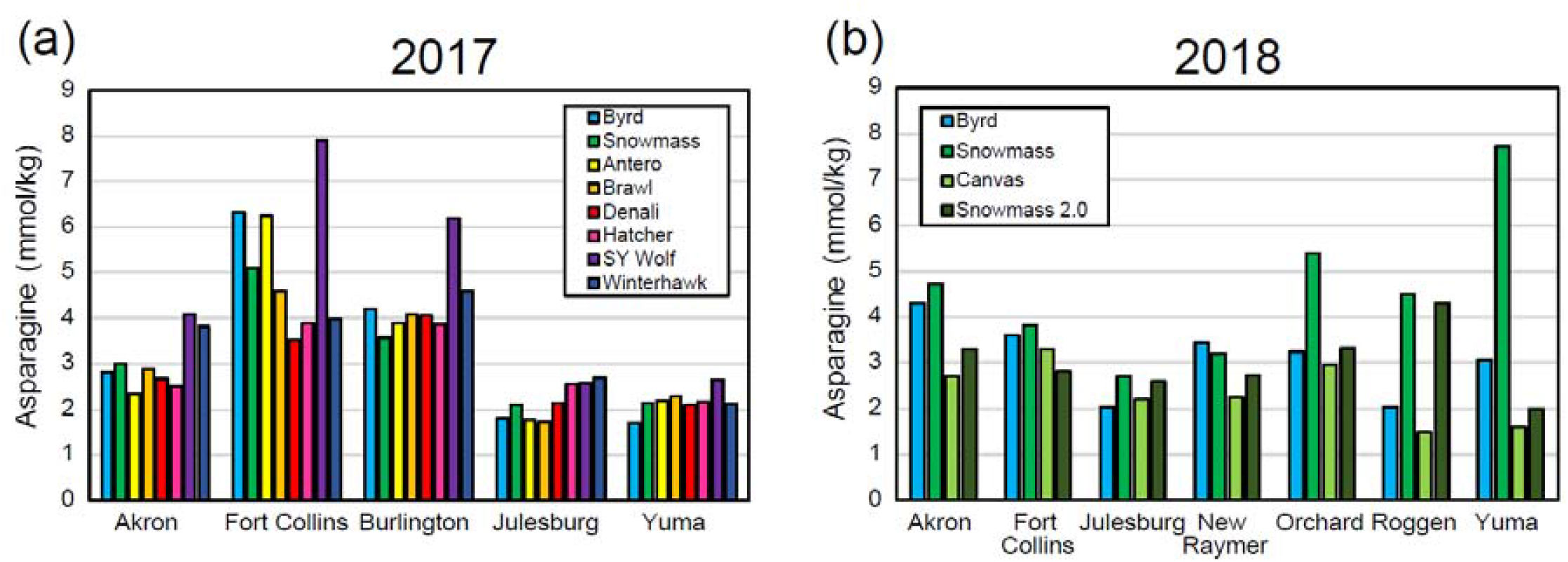
Free asparagine concentrations in elite winter wheat cultivars in (**a)** 2017 and (**b)** 2018 field seasons. Each measurement was a single replicate from field-grown experiments.

## DISCUSSION

### Induced *asn-a2* mutations confer reduced free asparagine concentration

Although natural genetic variation for free asparagine concentration exists in wheat germplasm collections, this is a highly quantitative trait determined by multiple small-effect QTL, complicating the identification, characterization and deployment of natural variants (Corol et al., 2016; Emebiri, 2014; Rapp et al., 2018). As a complementary approach, *in silico* databases of chemically-mutagenized wheat populations provide rapid access to novel genetic variation (Krasileva et al., 2017) that can be deployed in breeding programs without regulation, for example, to develop high-amylose wheat varieties (Hazard et al., 2012). One drawback of mutagenized populations is the confounding effect of residual background mutations, since tetraploid and hexaploid M_4_ plants carry, on average, 2,705 and 5,351 exonic point mutations, respectively (Krasileva et al., 2017). Even though two backcrosses were performed in the current study, the variable effect of these background mutations may account for phenotypic variation in line T4-2032 such that the differences in free asparagine concentration between wild-type and mutant sister lines were not significant (Figure 2). Despite this variation, consistent reductions in free asparagine concentrations in three independently-derived mutant populations in both tetraploid and hexaploid wheat provide strong evidence that the observed phenotype is conferred by non-functional induced mutations in *ASN-A2*. Furthermore, transcript levels of *ASN-B2* and *ASN-D2* were not affected by *asn-a2* mutations in any line (Figure 1), suggesting the impacts are due to reduced activity of *ASN-A2* itself, and not secondary effects on the activity of homeologous *ASN2* genes.

The reduction in free asparagine concentration in the *asn-a2* mutants ranged from 9% to 34% (Figure 2), comparable to the 16.2% reduction associated with a natural deletion of *TaASN-B2* in S-sufficient conditions (Oddy et al., 2021), but much lower than the 90% reduction observed in one CRISPR/Cas edited plant carrying mutations in all three *ASN2* homeologues (Raffan et al., 2021). The polyploid wheat genome provides functional redundancy, so it will be interesting to characterize isogenic materials with different combinations of *ASN2* alleles to reveal the relative impacts of each homeologue and the extent to which they act additively to reduce free asparagine concentration. These alleles may also be combined with independent natural QTL elsewhere in the genome to further lower free asparagine concentration (Emebiri, 2014; Rapp et al., 2018). It will be essential that these materials are tested in multi-environment yield plots including variation in S supply, and in more diverse genetic backgrounds in order to fully characterize the impact of environmental and G × E interactions for asparagine concentration (Emebiri, 2014; Rapp et al., 2018). Similar field trials using isogenic materials carrying higher-order combinations of *ASN2* alleles will reveal the extent to which it is possible to engineer reduced free asparagine concentration without introducing detrimental effects on agronomic or quality traits.

When integrating novel genetic variation, it is essential to determine possible pleiotropic effects on other traits. Most notably, CRISPR-edited plants with the greatest reductions in free asparagine concentration exhibited poor germination rates (Raffan et al., 2021), which may limit their utility for wheat breeders and growers. By contrast, the *asn-a2* mutant lines described in the current study exhibit no significant reduction in germination rate (Table 1), suggesting that the asparagine concentration in these seeds is sufficient for normal germination. A comprehensive quality analysis including SRC, flour and mixograph analyses revealed that *asn-a2* mutant lines exhibit no detrimental effects for quality traits compared to wild-type plants (Table 2). These findings are consistent with previous analyses of the impacts of natural variation in free asparagine concentration on these traits, where only mild negative correlations between sedimentation volume (a proxy for the quality traits measured here) and free asparagine concentration (Corol et al., 2016; Emebiri, 2014; Rapp et al., 2018). These results are encouraging and support the hypothesis that it should be possible to breed for mild reductions in asparagine concentration without compromising on baking quality or yield (Rapp et al., 2018).

### Reducing free asparagine concentration in other species

These findings show the potential to apply induced or natural variation in *ASN* genes to reduce asparagine concentrations in other species. Characterizing the expression profiles and natural variation of *ASN2* orthologs in barley (Avila-Ospina et al., 2015) and rye (Raffan & Halford, 2021) could reveal opportunities to reduce free asparagine concentration in the grain of these species. However, because of their diploid genomes, the strategy to induce more subtle reductions in asparagine concentration by targeting different combinations of homeologous genes would not be possible, and instead, milder variants that reduce, but do not abolish, gene activity may be more valuable.

The duplication event that gave rise to *TaASN1* and *TaASN2* genes occurred only in the Triticeae lineage, meaning that the genomes of other monocots, including rice, maize and sorghum, have no *ASN2* ortholog (Raffan & Halford, 2021). The rice genome contains two asparagine synthetase genes, so induced variation in either of these genes is likely to confer major changes in plant fitness not observed in wheat due to functional redundancy. However, in potato, while suppressing expression of both *StASN1* and *StASN2* had detrimental effects on growth and development (Chawla et al., 2012; Rommens et al., 2008), a more targeted approach focused only on *StASN1* reduced tuber asparagine content without other undesirable phenotypes, showing that it would be worthwhile exploring genetic diversity in the *ASN* gene family in other species.

### Characterizing the *ASN* gene family in wheat

Within wheat, reverse genetics approaches using either TILLING or CRISPR/Cas can also be applied to characterize other members of the *TaASN* gene family. In addition to their functional characterization, this approach may reveal novel combinations of alleles that could be beneficial for acrylamide reduction in wheat. Of note are *TaASN3*.*1* genes that are expressed during early embryo, ovule and grain development (Oddy et al., 2021; Xu et al., 2017). It would be interesting to develop mutants for *ASN3*.*1* genes to determine their contribution to free asparagine levels in the wheat grain and it is possible that combinations of these alleles with *ASN2* mutants could deliver further reductions in free asparagine concentrations in the grain. However, *TaASN3*.*1* genes are also expressed in leaf and stem tissues, so knockout alleles may have pleotropic effects on plant health and development.

## Conclusions

Non-transgenic, induced genetic variants in *ASN-A2* genes confer significant reductions in free asparagine concentration in wheat grains with no detrimental effects on agronomic or quality traits in field conditions. These alleles can be utilized immediately and without restrictions in breeding programs to reduce the acrylamide-forming potential of common wheat, either through recombination with the lines developed in this study, or by direct, targeted mutagenesis in elite wheat varieties.

## Abbreviations

ASN: asparagine synthetase
ATP: adenosine triphosphate
CRISPR/Cas: Clustered Regularly Interspaced Short Palindromic Repeats/CRISPR-associated protein
CTAB: Cetyl TrimethylAmmonium Bromide
DAA: days after anthesis
EMS: Ethyl MethaneSulfonate
G × E: genotype by environment
GATase: Glutamine Amidotransferase
IS: internal standard
KASP: Kompetitive Allele Specific PCR
MS: mass spectrometry
MTBE: Methyl-Tert-Butyl Ether
NIRS: Near Infrared Spectroscopy
QC: quality control
QTL: quantitative trait locus
qRT-PCR: Quantitative Reverse Transcriptase PCR
RNAi: Ribonucleic acid interference
S: sulphur
SKCS: Single Kernel Characterization System
SRC: Solvent Retention Capacity
TGW: Thousand Grain Weight
UPLC: Ultra-Performance Liquid Chromatography
Z-SDS: Z-Sodium Dodecyl Sulfate

## Supplemental Material

A PDF file containing Supplemental Figures S1, S2, S3, S4 and S5, and Supplemental Table S1.

## Conflicts of interest

The authors state no conflicts of interest.

## Author contributions

RAR – Investigation, formal analysis; YX – Investigation, formal analysis; JS - Investigation; JC - Investigation; REM – Funding acquisition, supervision; SP – Conceptualization, writing – original draft.

## Acknowledgements

This study was supported by funding from Colorado Wheat Research Foundation and the Colorado Wheat Administrative Committee. We are grateful to Forrest Wold-McGimsey for excellent technical support, to Dr. Jorge Dubcovsky for providing EMS-mutant lines for this study, and to Dr. Scott Haley for providing seeds from winter wheat elite lines grown in CSU winter wheat breeding plots. Asparagine quantification was performed at Colorado State University, by Dr. Linxing Yao, of the Analytical Resources Core: Bioanalysis and Omics Center (ARC-BIO).

## Notes

### Competing Interest Statement

The authors have declared no competing interest.

## References

AACC. (2000). Approved methods of the American Association of Cereal Chemists (AACC, Ed. 10th ed.).

Avila-Ospina, L., Marmagne, A., Talbotec, J., Krupinska, K., & Masclaux-Daubresse, C. (2015). The identification of new cytosolic glutamine synthetase and asparagine synthetase genes in barley (Hordeum vulgare L.), and their expression during leaf senescence. J Exp Bot, 66(7), 2013–2026. https://doi.org/10.1093/jxb/erv003

Beland, F. A., Mellick, P. W., Olson, G. R., Mendoza, M. C., Marques, M. M., & Doerge, D. R. (2013). Carcinogenicity of acrylamide in B6C3F(1) mice and F344/N rats from a 2-year drinking water exposure. Food Chem Toxicol, 51, 149–159. https://doi.org/10.1016/j.fct.2012.09.017

Bettge, A. D., Morris, C. F., DeMacon, V. L., & Kidwell, K. K. (2002). Adaptation of AACC method 56-11, solvent Retention capacity, for use as an early generation selection tool for cultivar development. Cereal Chemistry, 79, 670–674.

Bloksma, A. H., & Bushuk, W. (1988). Rheology and chemistry of dough. In Wheat chemistry and technology (3rd ed., pp. 131–218).

Chang, S., Puryear, J., & Cairney, J. (1993). A simple and efficient method for isolating RNA from pine trees. Plant Molecular Biology Reporter, 11(2), 113–116. https://doi.org/10.1007/BF02670468

Chawla, R., Shakya, R., & Rommens, C. M. (2012). Tuber-specific silencing of asparagine synthetase-1 reduces the acrylamide-forming potential of potatoes grown in the field without affecting tuber shape and yield. Plant Biotechnol J, 10(8), 913–924. https://doi.org/10.1111/j.1467-7652.2012.00720.x

Claus, A., Carle, R., & Schieber, A. (2008). Acrylamide in cereal products: A review. Journal of Cereal Science, 47(2), 118–133. https://doi.org/https://doi.org/10.1016/j.jcs.2007.06.016

Claus, A., Schreiter, P., Weber, A., Graeff, S., Herrmann, W., Claupein, W., Schieber, A., & Carle, R. (2006). Influence of agronomic factors and extraction rate on the acrylamide contents in yeast-leavened breads. J Agric Food Chem, 54(23), 8968–8976. https://doi.org/10.1021/jf061936f

Corol, D. I., Ravel, C., Rakszegi, M., Charmet, G., Bedo, Z., Beale, M. H., Shewry, P. R., & Ward, J. L. (2016). (1)H-NMR screening for the high-throughput determination of genotype and environmental effects on the content of asparagine in wheat grain. Plant Biotechnol J, 14(1), 128–139. https://doi.org/10.1111/pbi.12364

Curtis, T. Y., Postles, J., & Halford, N. G. (2014). Reducing the potential for processing contaminant formation in cereal products. J Cereal Sci, 59(3), 382–392. https://doi.org/10.1016/j.jcs.2013.11.002

Curtis, T. Y., Raffan, S., Wan, Y., King, R., Gonzalez-Uriarte, A., & Halford, N. G. (2019). Contrasting gene expression patterns in grain of high and low asparagine wheat genotypes in response to sulphur supply. BMC Genomics, 20(1), 628. https://doi.org/10.1186/s12864-019-5991-8

Emebiri, L. C. (2014). Genetic variation and possible SNP markers for breeding wheat with low-grain asparagine, the major precursor for acrylamide formation in heat-processed products. J Sci Food Agric, 94(7), 1422–1429. https://doi.org/10.1002/jsfa.6434

Gaines, C. S., Finney, P. F., Fleege, L. M., & Andrews, L. (1996). Predicting a hardness measurement using the single-kernel characterization system. Cereal Chemistry, 73, 278–283.

Gaufichon, L., Reisdorf-Cren, M., Rothstein, S. J., Chardon, F., & Suzuki, A. (2010). Biological functions of asparagine synthetase in plants. Plant Science, 179(3), 141–153. https://doi.org/https://doi.org/10.1016/j.plantsci.2010.04.010

Hazard, B., Zhang, X., Colasuonno, P., Uauy, C., Beckles, D. M., & Dubcovsky, J. (2012). Induced mutations in the starch branching enzyme II (SBEII) genes increase amylose and resistant starch content in durum wheat. Crop science, 52(4), 1754–1766. https://doi.org/10.2135/cropsci2012.02.0126

IARC. (1994). Some industrial chemicals: International Agency for Research on Cancer monographs on the evaluation of carcinogenesis risks to humans. W. H. O. Lyon (FR). https://www.ncbi.nlm.nih.gov/books/NBK507457/

JASP. (2019). JASP (Version 0.11.1) [Computer software]. In https://jasp-stats.org/

Krasileva, K. V., Vasquez-Gross, H. A., Howell, T., Bailey, P., Paraiso, F., Clissold, L., Simmonds, J., Ramirez-Gonzalez, R. H., Wang, X., Borrill, P., Fosker, C., Ayling, S., Phillips, A. L., Uauy, C., & Dubcovsky, J. (2017). Uncovering hidden variation in polyploid wheat. Proc Natl Acad Sci U S A, 114(6), E913–e921. https://doi.org/10.1073/pnas.1619268114

Lea, P. J., Sodek, L., Parry, M. A. J., Shewry, P. R., & Halford, N. G. (2007). Asparagine in plants. Annals of Applied Biology, 150(1), 1–26. https://doi.org/10.1111/j.1744-7348.2006.00104.x

Mottram, D. S., Wedzicha, B. L., & Dodson, A. T. (2002). Acrylamide is formed in the Maillard reaction. Nature, 419(6906), 448–449. https://doi.org/10.1038/419448a

Murray, M. G., & Thompson, W. F. (1980). Rapid isolation of high molecular weight plant DNA. Nucleic acids research, 8(19), 4321–4325. https://doi.org/10.1093/nar/8.19.4321

Navrotskyi, S., Baenziger, P. S., Regassa, T., Guttieri, M. J., & Rose, D. J. (2018). Variation in asparagine concentration in Nebraska wheat. Cereal Chemistry, 95(2), 264–273. https://doi.org/10.1002/cche.10023

Oddy, J., Alarcón-Reverte, R., Wilkinson, M., Ravet, K., Raffan, S., Minter, A., Mead, A., Elmore, J. S., de Almeida, I. M., Cryer, N. C., Halford, N. G., & Pearce, S. (2021). Reduced free asparagine in wheat grain resulting from a natural deletion of TaASN-B2: investigating and exploiting diversity in the asparagine synthetase gene family to improve wheat quality. BMC plant biology, 21(1), 302–302. https://doi.org/10.1186/s12870-021-03058-7

Raffan, S., & Halford, N. G. (2019). Acrylamide in food: Progress in and prospects for genetic and agronomic solutions. Annals of Applied Biology, 175(3), 259–281. https://doi.org/10.1111/aab.12536

Raffan, S., & Halford, N. G. (2021). Cereal asparagine synthetase genes. Annals of Applied Biology, 178(1), 6–22. https://doi.org/https://doi.org/10.1111/aab.12632

Raffan, S., Sparks, C., Huttly, A., Hyde, L., Martignago, D., Mead, A., Hanley, S. J., Wilkinson, P. A., Barker, G., Edwards, K. J., Curtis, T. Y., Usher, S., Kosik, O., & Halford, N. G. (2021). Wheat with greatly reduced accumulation of free asparagine in the grain, produced by CRISPR/Cas9 editing of asparagine synthetase gene TaASN2. Plant Biotechnol J. https://doi.org/10.1111/pbi.13573

Rapp, M., Schwadorf, K., Leiser, W. L., Wurschüm, T., & Longin, C. F. H. (2018). Assessing the variation and genetic architecture of asparagine content in wheat: What can plant breeding contribute to a reduction in the acrylamide precursor? Theor Appl Genet, 131(11), 2427–2437. https://doi.org/10.1007/s00122-018-3163-x

Rommens, C. M., Yan, H., Swords, K., Richael, C., & Ye, J. (2008). Low-acrylamide French fries and potato chips. Plant biotechnology journal, 6(8), 843–853. https://doi.org/10.1111/j.1467-7652.2008.00363.x

Stadler, R. H., Blank, I., Varga, N., Robert, F., Hau, J., Guy, P. A., Robert, M. C., & Riediker, S. (2002). Acrylamide from Maillard reaction products. Nature, 419(6906), 449–450. https://doi.org/10.1038/419449a

Surdyk, N., Rosen, J., Andersson, R., & Aman, P. (2004). Effects of asparagine, fructose, and baking conditions on acrylamide content in yeast-leavened wheat bread. J Agric Food Chem, 52(7), 2047–2051. https://doi.org/10.1021/jf034999w

Tareke, E., Rydberg, P., Karlsson, P., Eriksson, S., & Törnqvist, M. (2002). Analysis of acrylamide, a carcinogen formed in heated foodstuffs. J Agric Food Chem, 50(17), 4998–5006. https://doi.org/10.1021/jf020302f

Wilson, T. L., Guttieri, M. J., Nelson, N. O., Fritz, A., & Tilley, M. (2020). Nitrogen and sulfur effects on hard winter wheat quality and asparagine concentration. Journal of Cereal Science, 93, 102969. https://doi.org/https://doi.org/10.1016/j.jcs.2020.102969

Xu, H., Curtis, T. Y., Powers, S. J., Raffan, S., Gao, R., Huang, J., Heiner, M., Gilbert, D. R., & Halford, N. G. (2017). Genomic, biochemical, and modeling analyses of asparagine synthetases from wheat. Front Plant Sci, 8, 2237. https://doi.org/10.3389/fpls.2017.02237

Zhivagui, M., Ng, A. W. T., Ardin, M., Churchwell, M. I., Pandey, M., Renard, C., Villar, S., Cahais, V., Robitaille, A., Bouaoun, L., Heguy, A., Guyton, K. Z., Stampfer, M. R., McKay, J., Hollstein, M., Olivier, M., Rozen, S. G., Beland, F. A., Korenjak, M., & Zavadil, J. (2019). Experimental and pan-cancer genome analyses reveal widespread contribution of acrylamide exposure to carcinogenesis in humans. Genome Res, 29(4), 521–531. https://doi.org/10.1101/gr.242453.118

