## Supplemental_materials for "INDUCED MUTATIONS IN *TaASN-A2* REDUCE FREE ASPARAGINE CONCENTRATION IN THE WHEAT GRAIN"

**Supplemental Figures S1, S2, S3, S4 and S5.**

**Supplemental Table S1.**

**Supplemental Figure S1:** KASP genotyping assay to detect G468A mutation in line T4-1388. **(a)** Position of primers in *ASN-A2* target gene. **(b)** Representative plot of expected genotypic classes.

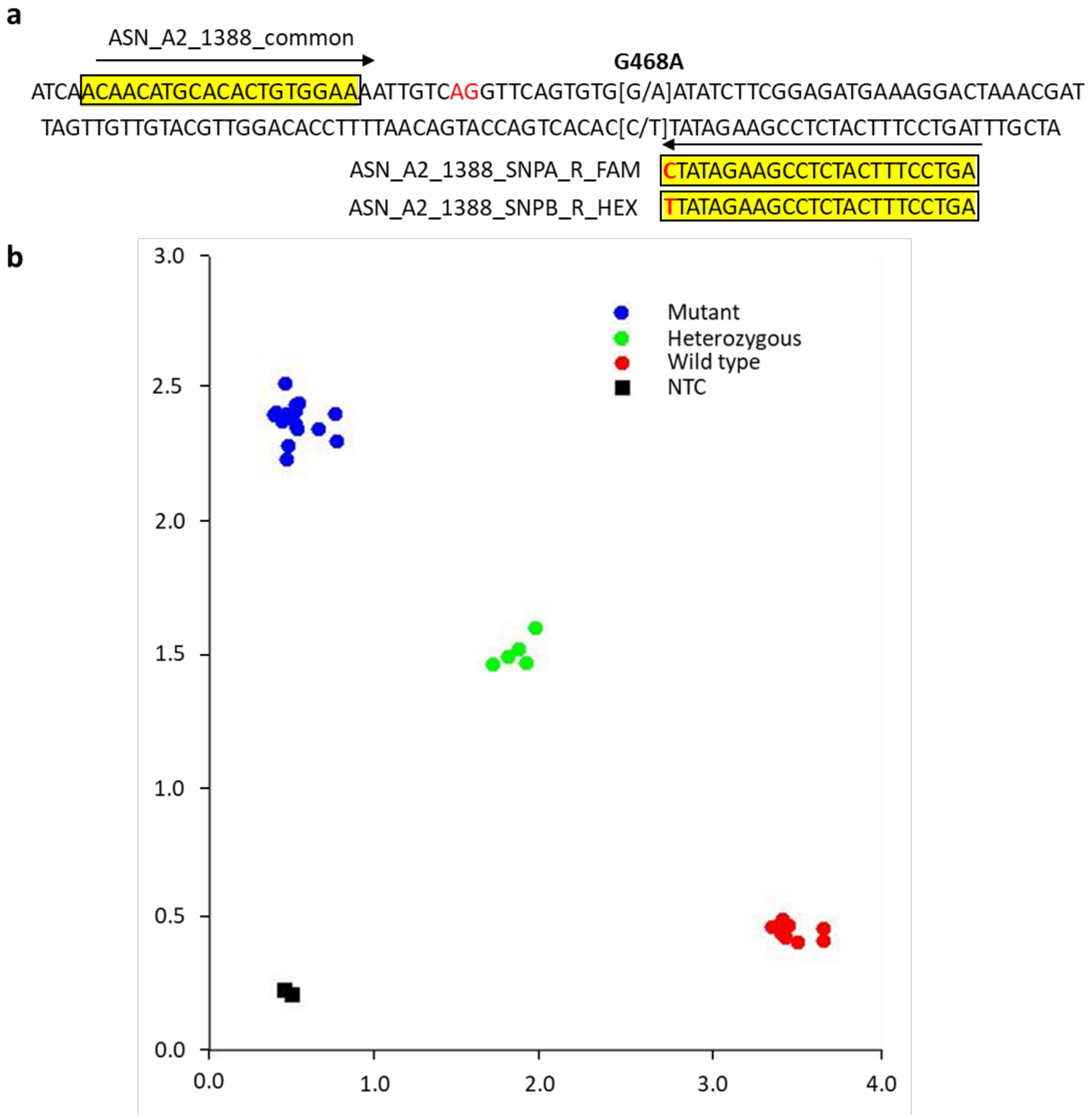

**Supplemental Figure S2:** KASP genotyping assay to detect G446A mutation in line T4-2032. **(a)** Position of primers in *ASN-A2* target gene. **(b)** Representative plot of expected genotypic classes.

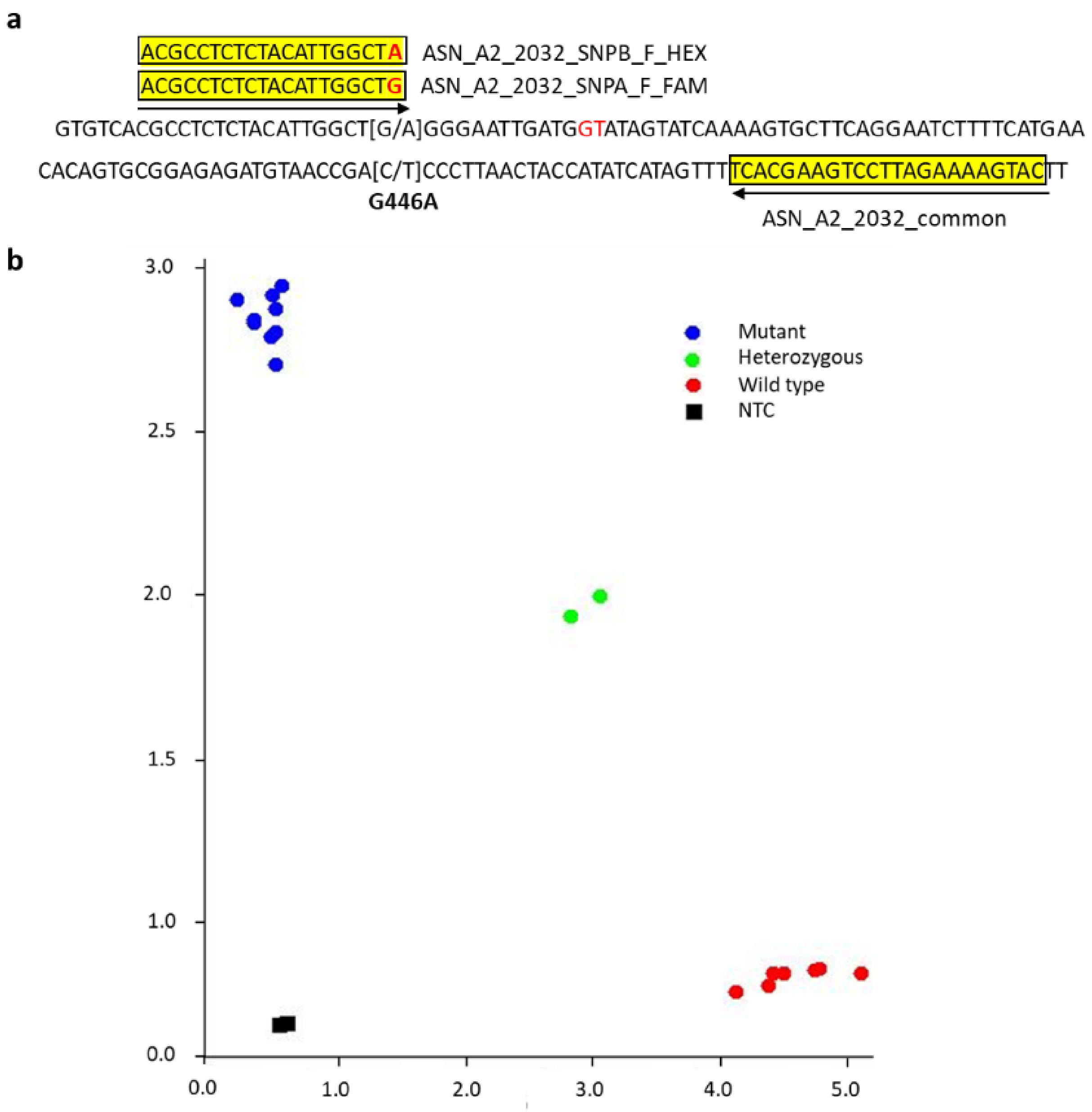

**Supplemental Figure S3:** CAPS marker to detect mutation G585A in line T6-1048. PCR products were digested with *StyI*. Template DNA from plants carrying the A residue at this position (Mutant type, MT) were not digested and present as a 1,063bp product. Template DNA from plants carrying the G residue at this position (Wild-type, WT) were digested into two products, 946 bp and 117 bp (not visible on this gel). Heterozygous plants exhibit a mix of both products (Het).

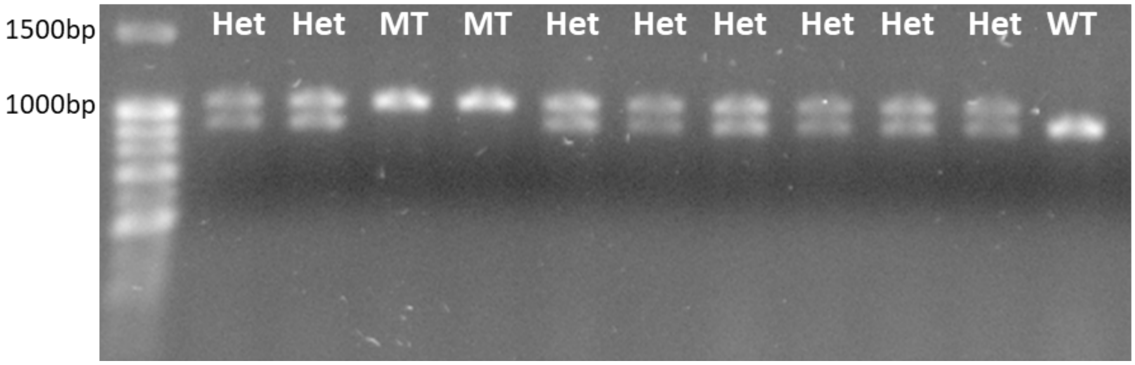

**Supplemental Figure S4:** PCR assay to distinguish presence and absence of *TaASN-B2* iSn a collection of ten elite winter wheat cultivars and three control lines (Cadenza, Kronos and Chinese Spring). **a.** Schematic diagram of the assay to show primer positions and expected amplicon sizes. Amplification of a 189 bp product with primers P1 and P2 indicates that *TaASN-B2* is deleted, while amplification of a 125 bp product with primers P3 and P4 indicates that *TaASN-B2* is present. One amplified fragment is expected in each reaction. **b.** Agarose gel electrophoresis of PCR products from the assay. Varieties with *TaASN-B2* deleted are highlighted in red, while varieties with the gene present are highlighted in green. A 100 bp ladder is shown in the first and last well of the gel for size comparison.

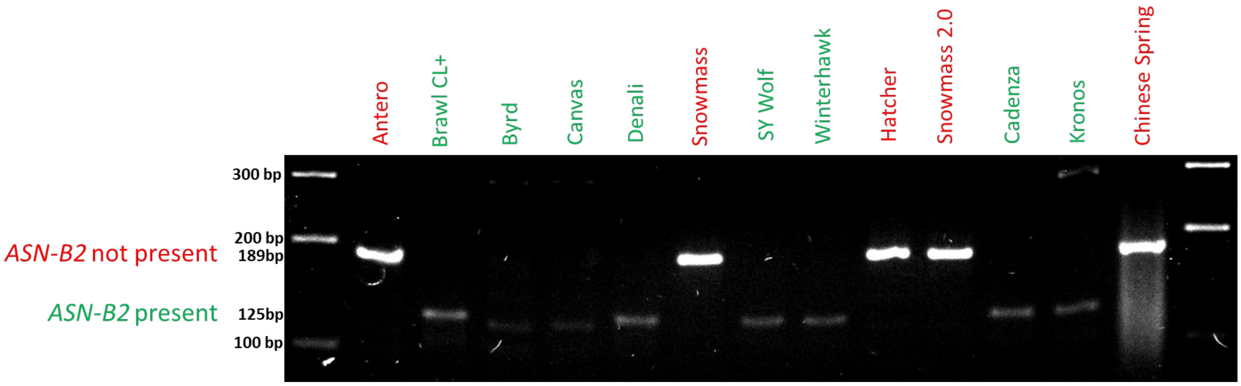

**Supplemental Figure S5**: Raw mixograph curves for each sample, separated by population and genotype.

#### T6-1048 wild-type

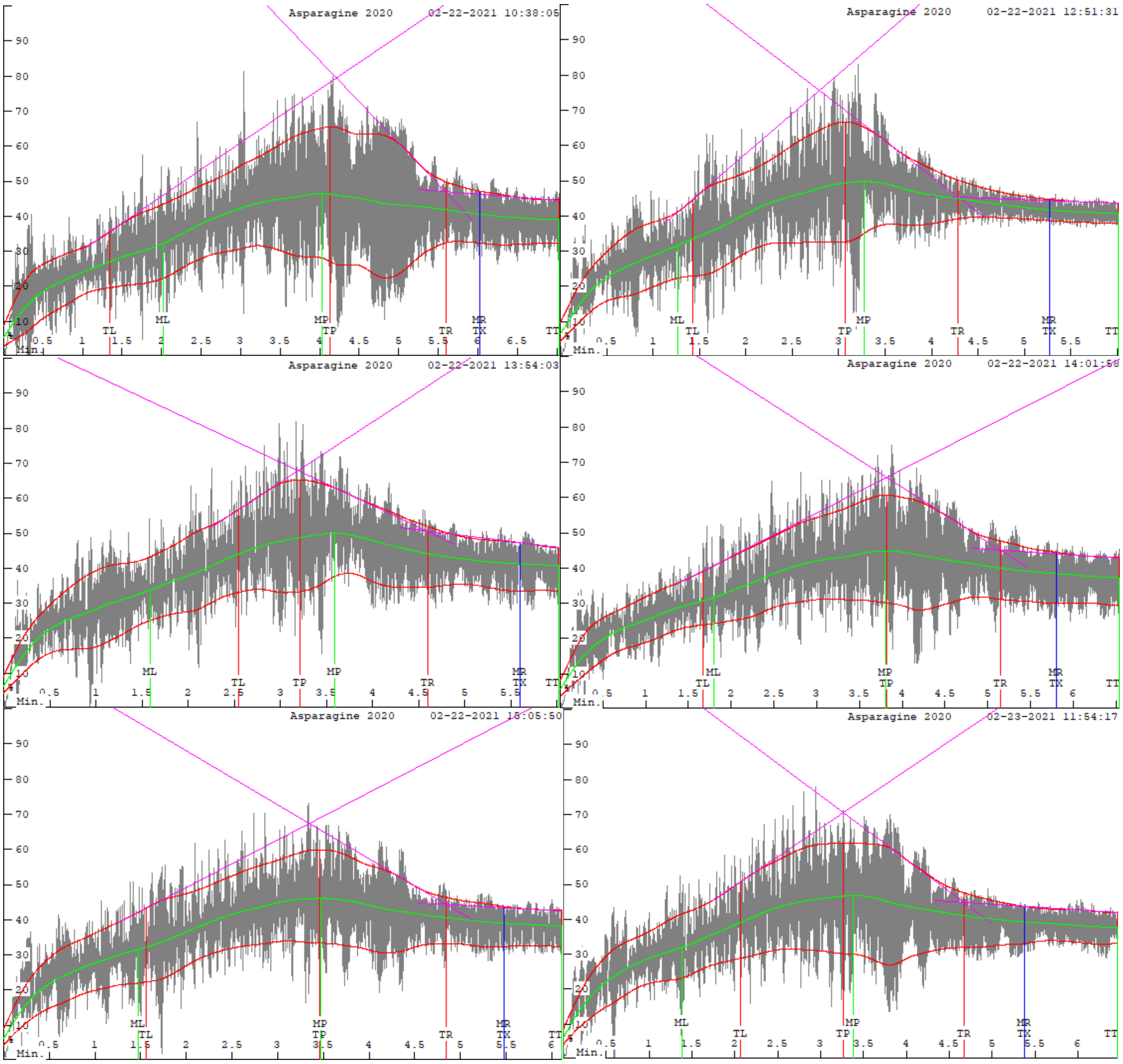

**T6-1048 *asn-a2***

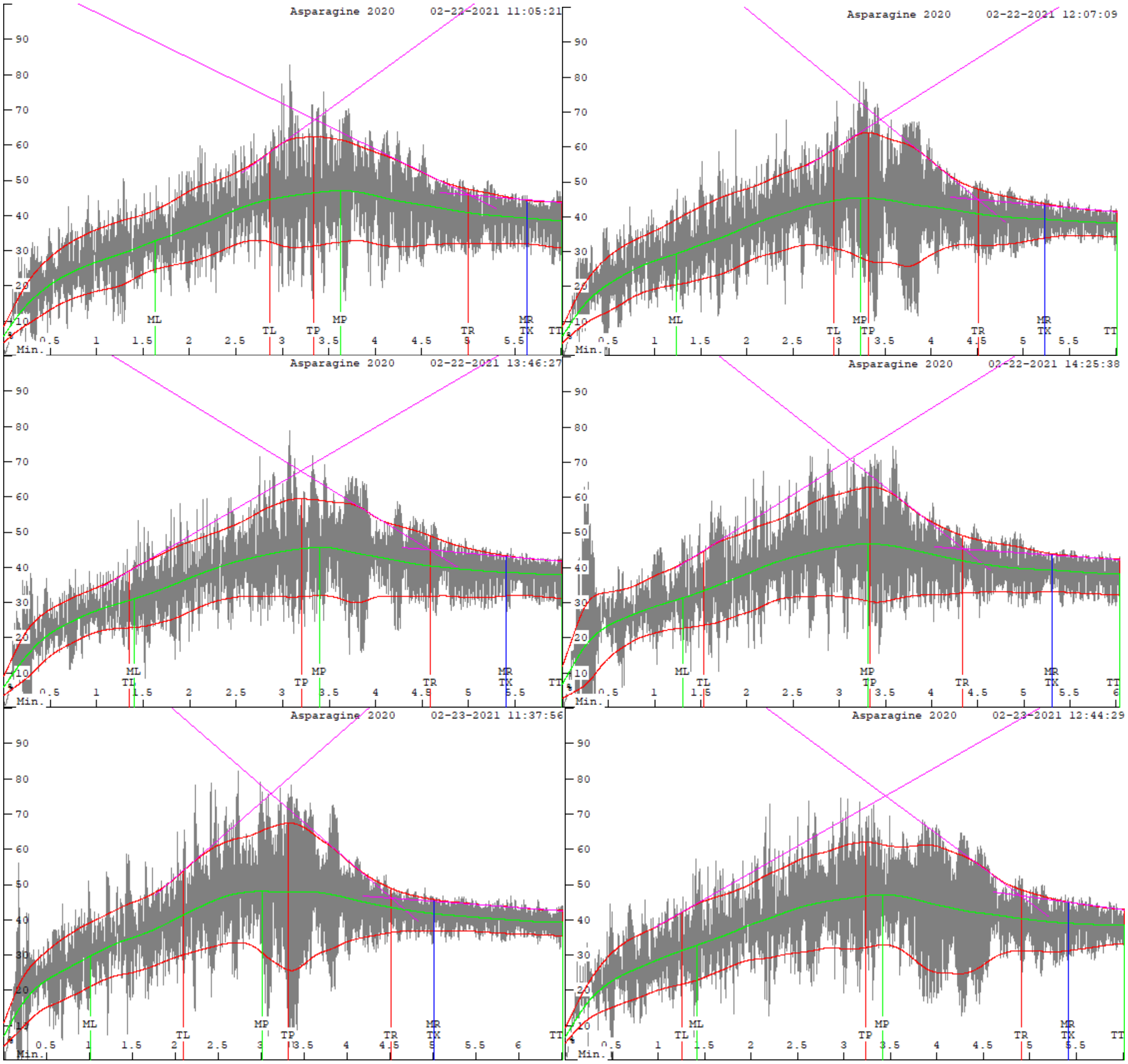

### T4-1388 wild-type

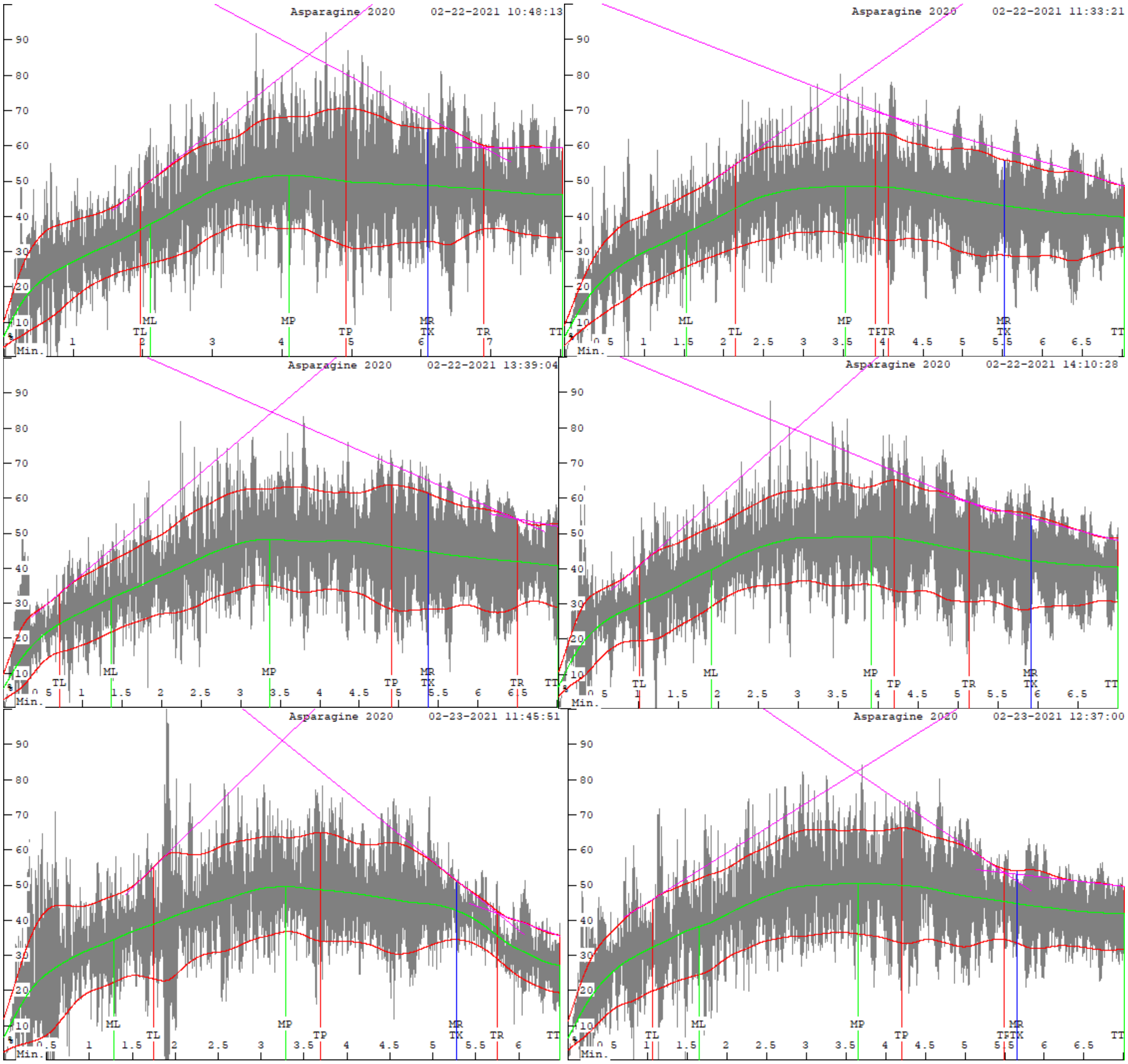

### T4-1388 *asn-a2*

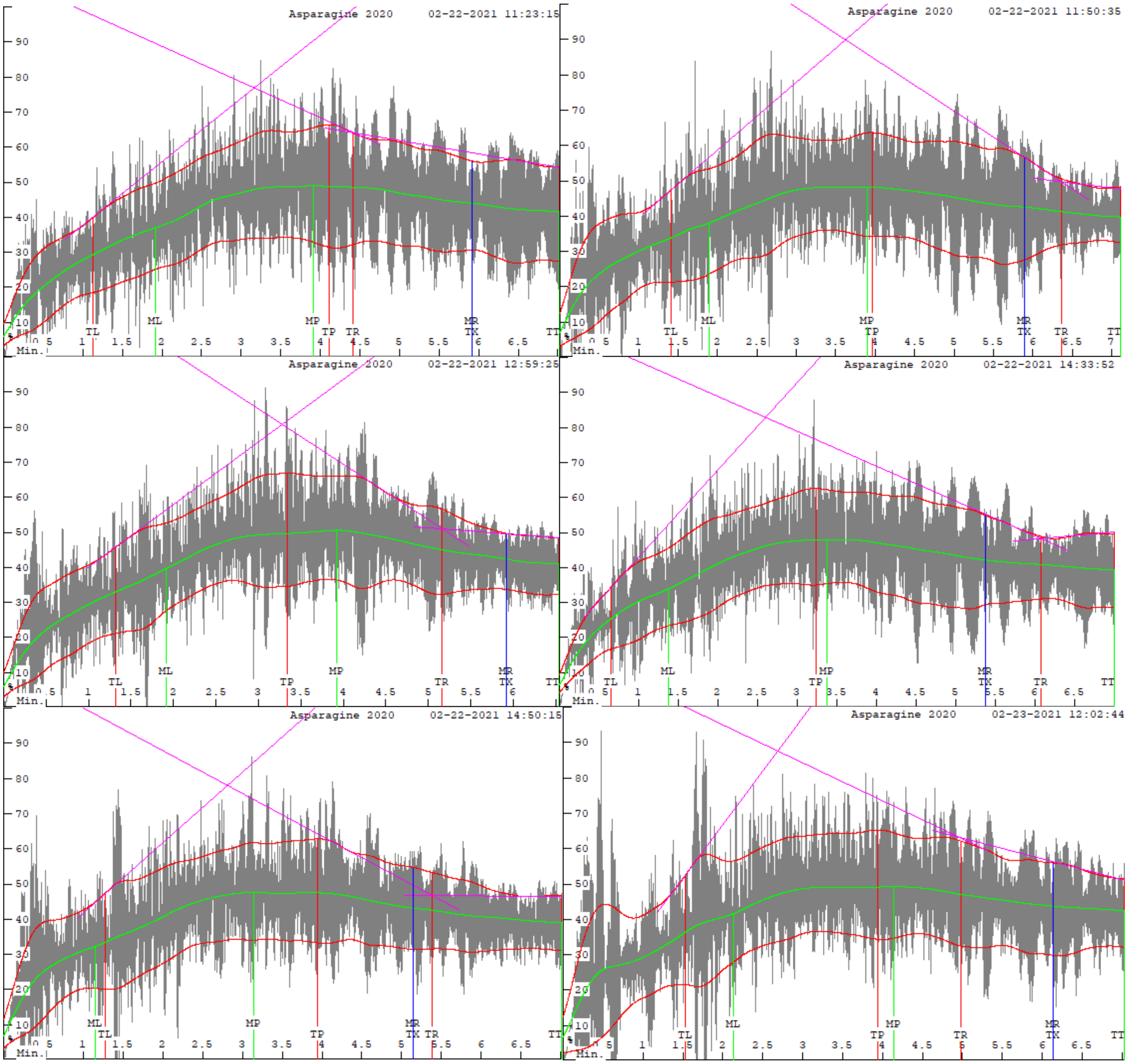

### T4-2032 wild-type

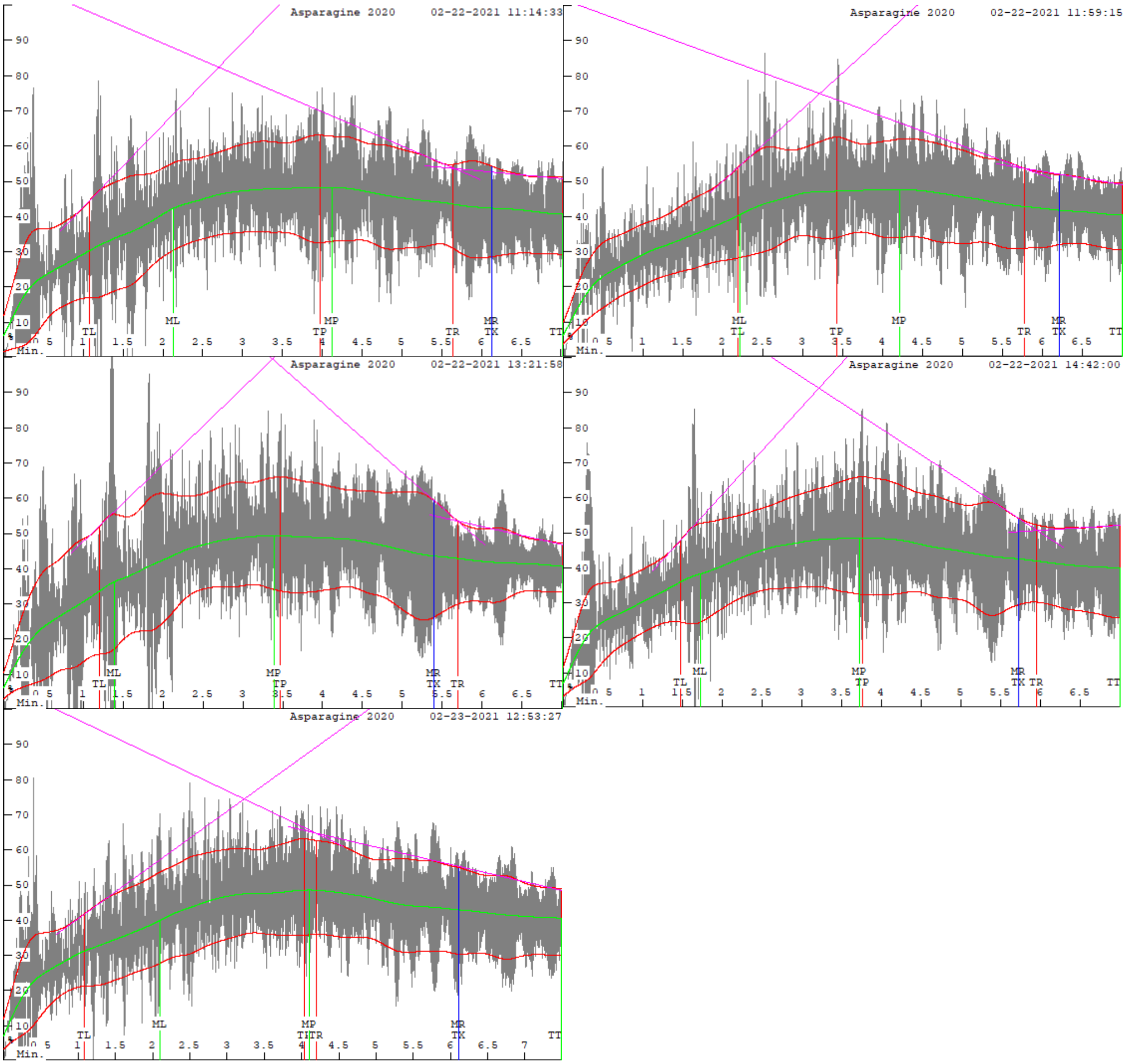

### T4-2032 *asn-a2*

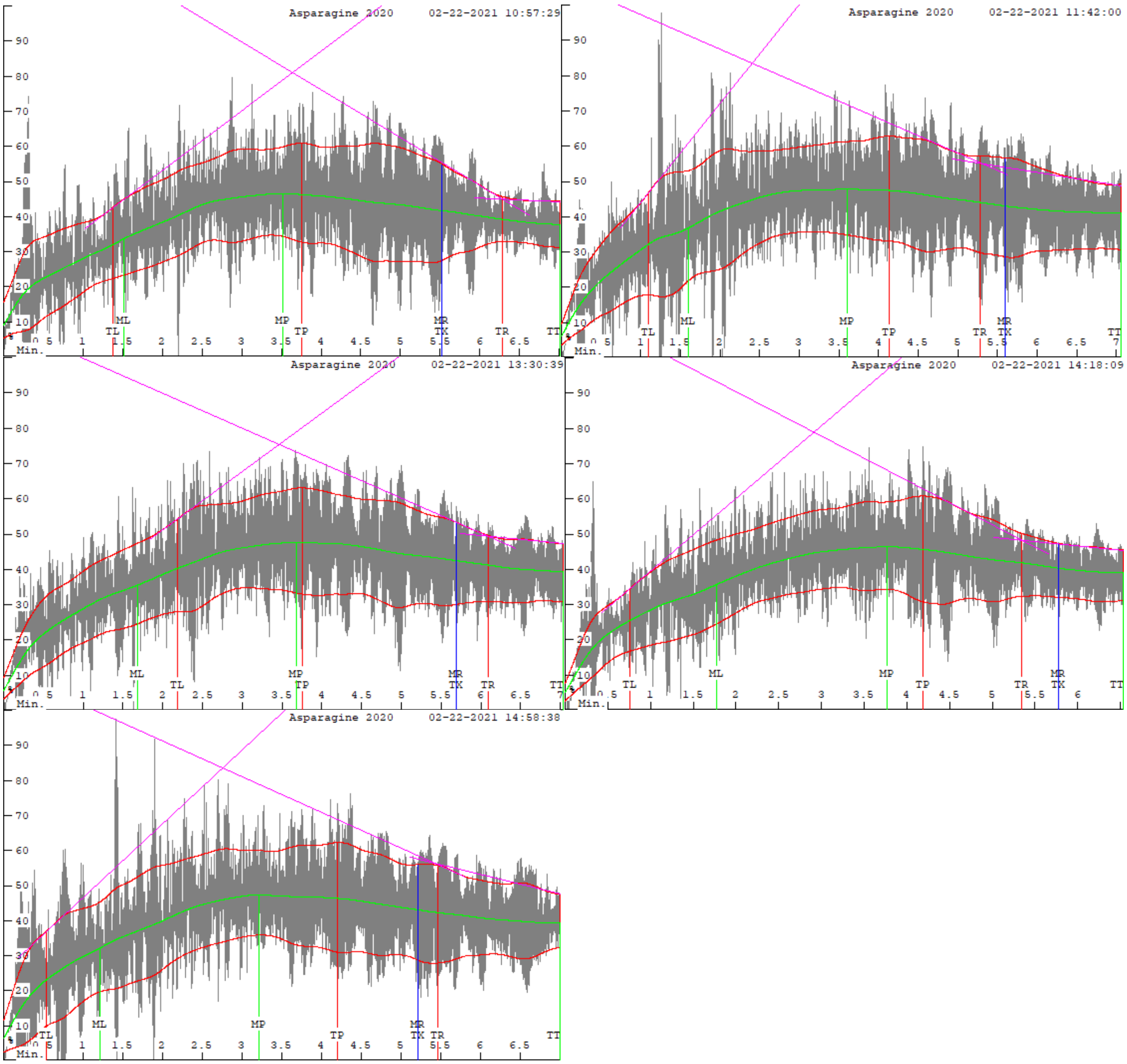

**Supplemental Table S1:** Primers used in this study.

| Purpose | Amplified region | Primer Name | Primer Sequence (5'-3') | Amplicon size (bp) |
|---|---|---|---|---|
| Check for the absence of ASN-B2 | Region flanking deleted region in varieties with no ASN-B2 | ASN-B2_CS_F3 | AGCAAGCCTTCACCATCATT | 189 |
| | | ASN-B2_CS_R1 | GATGTAGGCATGTCAACGAGA | |
| Check for the presence of ASN-B2 | Exon 11-3' UTR | ASN-B2_qF1 | AACAAGCCTGGGGTGATGAG | 125 |
| | | ASN-B2_qR1 | TTGTCTCAAAAAGAAAAAGAACTTG | |
| T6-1048 CAPS marker | Region from intron 1 to 3 | ASN-A2_1048_F3 | AAGAAGAAAGTGACCACACA | 1,063 |
| | | ASN-A2_1048_R3 | TTTTCAAAGATACTAGCACATAAT | |
| T4-1388 KASP marker | Region from intron 2 to exon 3 | ASN_A2_1388_SNPA_R_FAM | GAAGGTGACCAAGTTCATGCTAGTCCTTTCATCT CCGAAGATATC | 62 |
| | | ASN_A2_1388_SNPB_R_HEX | GAAGGTCGGAGTCAACGGATTAGTCCTTTCATCT CCGAAGATATT | |
| | | ASN_A2_1388_common | ACAACATGCACACTGTGGAA | |
| T4-2032 KASP marker | Region from exon 2 to intron 2 | ASN_A2_2032_SNPA_F_FAM | GAAGGTGACCAAGTTCATGCTACGCCTCTCTACA TTGGCTG | 66 |
| | | ASN_A2_2032_SNPB_F_HEX | GAAGGTCGGAGTCAACGGATTACGCCTCTCTACA TTGGCTA | |
| | | ASN_A2_2032_common | CATGAAAAGATTCCTGAAGCACT | |
| *ASN-A2* expression | Exons 1 and 2 | ASN-A2_F | TCAACGGGGAGGTCTACAAC | 184 |
| | | ASN-A2_R | GCAATGAAGCTGTTATCTCGTG | |
| *ASN-B2* expression | Exons 1 and 2 | ASN-B2_F | GTCAACGGGGAGATCTACAACC | 185 |
| | | ASN-B2_R | GCGATGAAGCTGTGATCTCTTG | |
| *ASN-D2* expression | Exons 1 and 2 | ASN-D2_F | GTGAACGGGGAGATTTACAAC | 185 |
| | | ASN-D2_R | GCAATGAAGCTCTTATCTCGTG | |
| Housekeeping gene | | ACTIN-F | CAGAGTCGAGCACAATACCAGTTG | 91 |
| | | ACTIN-R | ACCTTCAGTTGCCCAGCAAT | |
